# *Spodoptera frugiperda* transcriptional response to infestation by *Steinernema carpocapsae*

**DOI:** 10.1101/618165

**Authors:** Louise Huot, Simon George, Pierre-Alain Girard, Dany Severac, Nicolas Nègre, Bernard Duvic

## Abstract

*Steinernema carpocapsae* is an entomopathogenic nematode (EPN) used in biological control of agricultural pest insects. It enters the hemocoel of its host via the intestinal tract and releases its symbiotic bacterium *Xenorhabdus nematophila*, which kills the insect in less than 48 hours. Although several aspects of its interactions with insects have been extensively studied, still little is known about the immune and physiological responses of its different hosts. In order to improve this knowledge, we examined the transcriptional responses to EPN infestation of the fat body, the hemocytes and the midgut in the lepidopteran pest model *Spodoptera frugiperda* (Lepidoptera: Noctuidae).

Our results indicate that the tissues poorly respond to the infestation at an early time post-infestation of 8 h, even though the proliferation of the bacterial symbiont within the hemocoel is detected. Only 5 genes are differentially expressed in the fat body of the caterpillars. However, strong transcriptional responses are observed at a later time point of 15 h post-infestation in all three tissues. While few genes are differentially expressed in the midgut, tissue-specific panels of induced metalloprotease inhibitors, immune receptors and antimicrobial peptides together with several uncharacterized genes are up-regulated in the fat body and the hemocytes. In addition, among the most up-regulated genes, we identified new potential immune effectors, unique to Lepidoptera, for which we present evidence of acquisition by Horizontal Gene Transfer from bacteria.

Altogether, these results pave the way for further functional studies of the mobilized genes’ involvement in the interaction with the EPN.

**Author summary:** The Fall Armyworm, *Spodoptera frugiperda*, is a major agricultural pest. The caterpillars cause extensive damage to crops of importance such as corn, rice, sorghum and cotton. Originally from the Americas, it is currently becoming invasive in other parts of the world, first in Africa in 2016, then in India and now in south-east Asia. Programs of biological control against insect pests are increasingly encouraged around the world and include the use of pathogens. Entomopathogenic nematodes such as *Steinernema carpocapsae* are already commercialized as organic pesticides. These nematodes live in the soil and enter the body of their insect preys. Once within the insects, they release their symbiotic bacteria (*Xenorhabdus nematophila* in this case), which infect and kill the host in a few hours. The nematodes can then feed on the dead insects, reproduce and resume their life cycle. It is a major challenge to understand how EPN achieve their pathogenicity as well as how the insects can resist them. Here we provide the foundation for such an interaction between EPN and a Lepidoptera. We analyzed the dynamic of transcriptional response in three insect tissues (midgut, fat body and hemocytes) upon infestation by EPN. Not many studies have been performed genome-wide on such an interaction, and none on a Lepidopteran model of economical importance. Our transcriptomic approach revealed some specificities of the Lepidopteran defense mechanisms. In particular, we discovered a set of genes, acquired in Lepidoptera from bacteria by Horizontal Gene Transfer, that probably encode proteins with antibiotic activity.

## Introduction

There is a growing desire in Europe to reduce the use of chemical pesticides on agricultural land, because of their toxicity for the environment and human health (European Directive EC91/414). Also, the development of alternative methods for the control of crop pests is encouraged. These methods include the use of predators and pathogens of insect pests such as viruses, bacteria, fungi, parasitoid wasps and entomopathogenic nematodes (EPN).

EPN of the genus *Steinernema* associated with the symbiotic bacterium *Xenorhabdus* are among the most widely used and studied biological control agents (1). They pose little threat to human health and non-target species (2), but are capable of killing a broad spectrum of insect pests including the moth *Spodoptera frugiperda* (Lepidoptera: Noctuidae)(3, 4). The infestation cycle of the nematode *Steinernema carpocapsae* has been well described. It starts with the entry of infective juvenile larvae (IJ) into the insect intestinal tract via the natural orifices (5). Once the intestinal epithelium is crossed, EPN are found in the hemocoel where they release their symbiotic bacteria *Xenorhabdus nematophila*. The latter cause the death of the insect by toxemia and sepsis in less than 48 h. EPN multiply by feeding on the insect tissues, reassociate with their symbiotic bacteria (6) and then go back to the environment in search for a new prey (7).

Insects live in environments contaminated by microorganisms (bacteria, fungi or viruses) potentially pathogenic for them. To defend themselves against these aggressors, insects have developed a powerful and diversified immune system essentially based on innate immunity and which has been well described in the *Drosophila* model (8). Its main mechanisms and pathways seem to be conserved in different orders of insects (9–11) even though some insects, such as *Apis mellifera* or *Acyrthosiphon pisum*, have reduced immune repertoire (12, 13). The first line of defense are physical barriers, including the cuticle, which covers almost all insects’ interfaces with the environment, and the peritrophic matrix, which replaces the cuticle in the midgut (14). The cuticle is a thick exoskeleton made of wax, chitin and sclerotized proteins that confers mechanical protection against wounds and invaders (15), while the midgut peritrophic matrix is a thinner network of chitin and proteins that allows the uptake of nutrients but which has a low permeability to microorganisms and toxins (16). The intestinal epithelium can produce several immune molecules such as antimicrobial peptides (AMP) or reactive oxygen species depending on the location (14). Once into the hemolymph, parasites are facing circulating hemocytes, which are the immune blood cells of insects. Hemocytes participate in sclerotization, coagulation as well as in the elimination of small pathogens (bacteria and yeasts) by phagocytosis and nodulation, and of large pathogens (parasitoid wasp eggs, nematodes) by encapsulation (17). Encapsulation, nodulation and coagulation may involve a process of melanisation, which results in the formation of melanin and toxic chemicals that help to sequester and to kill the pathogen (18). Melanisation is activated by an extracellular proteolytic cascade, the pro-phenoloxidase system, following the recognition of microbial determinants or danger signals (19). Finally, the systemic response of insects relies mainly on the massive secretion of AMP by the fat body into the hemolymph, after activation of the IMD and/or Toll pathways. The IMD pathway is primarily activated by peptidoglycan recognition proteins (PGRP) in response to Gram-negative bacteria and the Toll pathway by PGRP and Gram negative binding proteins (GNBP) in response to Gram-positive bacteria and fungi (20). The Toll pathway may also be activated by exogenous proteases from pathogens (21).

Recently, different studies performed in the model insect, *Drosophila melanogaster*, described the responses of this insect to two EPN, *Steinernema carpocapsae* or *Heterorhabditis bacteriophora* (22–24). By the use of transcriptomic approaches on whole larvae or adult flies, the authors showed the overexpression of a large number of immune-related genes involved in defense responses such as coagulation, melanisation and the production of antimicrobial peptides, and in several immune and stress-reponse pathways (Toll, Imd, Jak/Stat or JNK).

In this study, we carried out a transcriptomic analysis of physiological and immune responses of 6^th^ instar larvae of *S. frugiperda* fat body, hemocytes and midgut at two time points after infection with the EPN *S. carpocapsae*. At 8 hours after infestation (hpi), we found only 5 genes that were differentially expressed in the fat body. However, at 15 hpi, we detected the significant expression modulation of 271 genes. Few genes were differentially expressed in the midgut whereas strong transcriptional responses were observed in the fat body and the hemocytes. These responses consisted mainly in the overexpression of induced metalloprotease inhibitors (IMPI), immune receptors (mostly PGRP) and AMP, indicating that the fat body as well as the hemocytes produce potent immune responses. Among the most up-regulated genes, we identified a cluster of new potential immune effectors, unique to Lepidoptera, for which we present evidence of acquisition by Horizontal Gene Transfer from bacteria. Finally, we identified a cluster of genes that are overexpressed in all tissues, but whose function is unknown and which are restricted to some noctuid species.

## Results and discussion

### EPN infestation and pathogenicity

To measure how *S. frugiperda* larvae respond to entomopathogenic nematodes (EPN), we performed infestation experiments where 6^th^ instar larvae were individually put in contact with either Ringer solution (control experiment) or a solution of EPN in Ringer (infestation condition) at time T = 0 (see Methods and Fig 1A). We targeted two time points after infestation based on our previous knowledge of the mode of infestation (6). Eight hours post infestation (hpi), nematodes are supposed to have travelled in the intestinal tract of *S. frugiperda* larvae, crossed the intestinal barrier and started releasing their symbiotic bacteria, *Xenorhabdus nematophila*, within the hemocoel of the caterpillar (6). At 15 hpi, bacteria have multiplied and septicemia is supposed to be reached. In order to verify these assertions, we quantified the *X. nematophila* cells into the hemolymph by CFU counting on a selective medium. In parallel, the survival of the caterpillar to the EPN infestation was monitored for 72 h. Our data show that bacterial growth has already started at 8 hpi, from few bacterial cells released to 10^4^ cells/mL of hemolymph and up to 10^6^ cells/mL at 15 hpi, which is considered septicemia (Fig 1B). When we measured survival of the larvae following infestation, we observed that the first deaths occur at 28 hpi. At 72 hpi, almost all treated larvae were dead (Fig 1C) whereas no death was observed in the control experiments.

**Fig 1:**
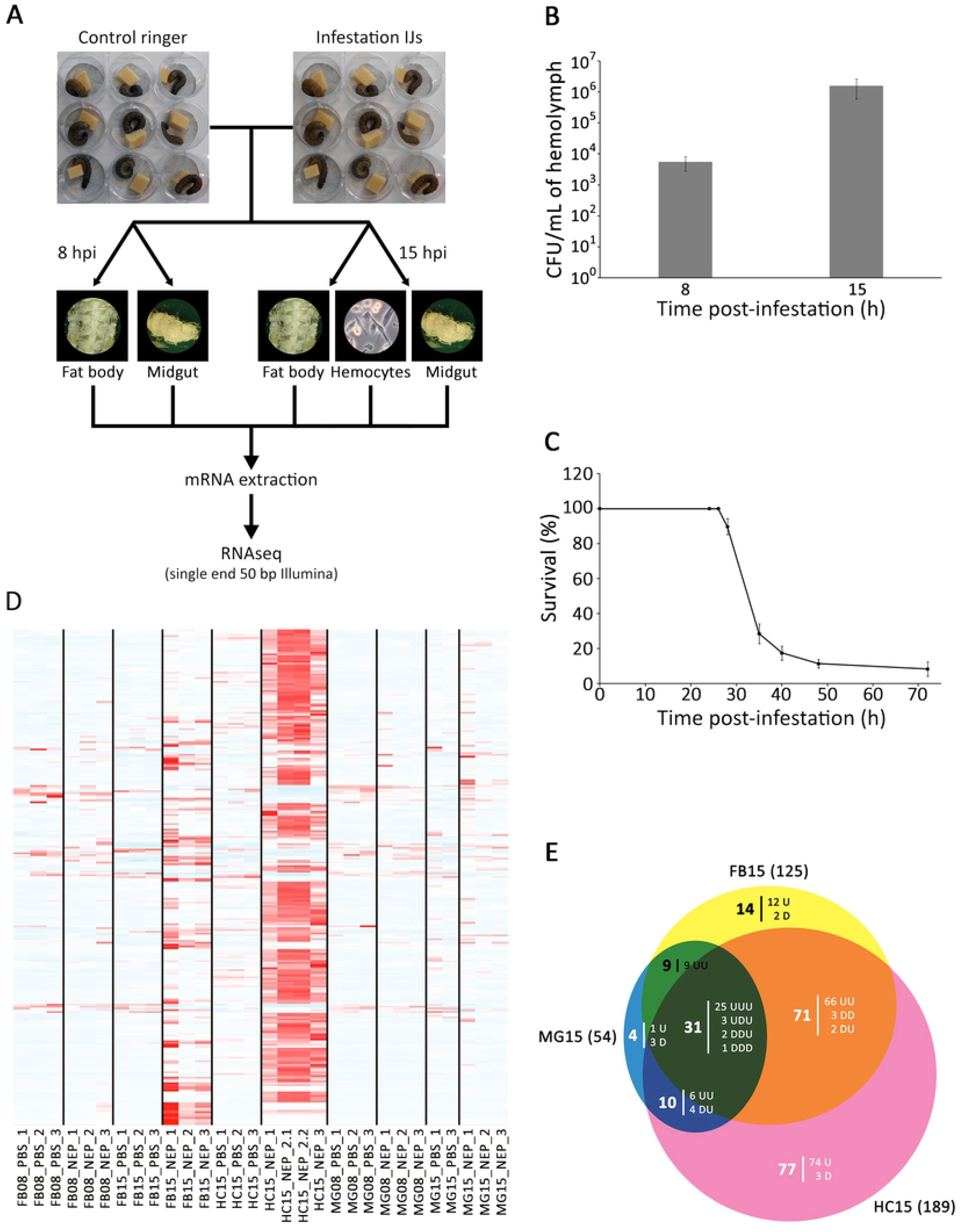
Tissue specific transcriptional response time series of *Spodoptera frugiperda* larvae to EPN infestation. **A:** Overview of the experimental design. In three independent experiments, 9 infested and 9 control larvae from culture plates were dissected at 8 hpi and 15 hpi. Hemocytes, fat bodies and midguts were extracted and pooled by organ for each time and condition. Polyadenylated RNAs were purified from these pools and corresponding cDNA libraries were built. cDNAs were sequenced on a single end by Illumina and RNAseq data were analyzed to identify the genes that are differentially expressed during *Steinernema carpocapsae* infestation. **B:** Growth of *Xenorhabdus nematophila* following *S. frugiperda* infestation with 150 symbiotic *S. carpocapsae*. At 8 hpi and 15 hpi, the number of CFU per mL of hemolymph was estimated from three independent experiments with three technical replicates (three larvae per technical replicate). Error bars indicate standard errors of the means. **C:** Survival curve. Larvae were infested with 150 nematobacterial IJ. Data represent means ± SEs of four independent experiments, each containing 12 larvae. **D:** EPN effect significant gene response. This heatmap shows z-score of expression variation across all RNAseq samples (red being overexpressed and blue under-expressed) for 271 genes with significant variations to EPN infestation. **E:** Venn diagram showing the tissue specificity of the EPN responsive genes at 15 hpi. The response can be overexpression (Up:U) or under-expression (Down:D). For example, there are 71 genes varying significantly upon EPN infestation in both the fat body (FB) and the hemocytes (HC). Of these, 66 are up-regulated in both tissues (UU), 3 are down-regulated in both tissues (DD) and 2 are down-regulated in FB and up-regulated in HC (DU). By convention, the order of the U and D letters represent respectively MG, FB and HC tissues.

At 8 hpi and 15 hpi, we removed the larvae from either the control or the infestation plates and dissected three tissues: the midgut (MG), the fat body (FB) and the hemocytes (HC at 15 hpi only). From these tissues, RNA was extracted and processed to perform single end 50 bp Illumina sequencing on three biological replicates (Fig 1A). We recovered between 10 and 100 million reads per sample, of which between 50 and 70% align onto the reference transcriptome for *S. frugiperda* (25) (see Methods and Table 1). After normalization with DESeq2 (26), we observed that gene expression datasets cluster by tissue (**S1 Fig**) and that within tissues, condition replicates correlate well with each other.

**Table 1:**
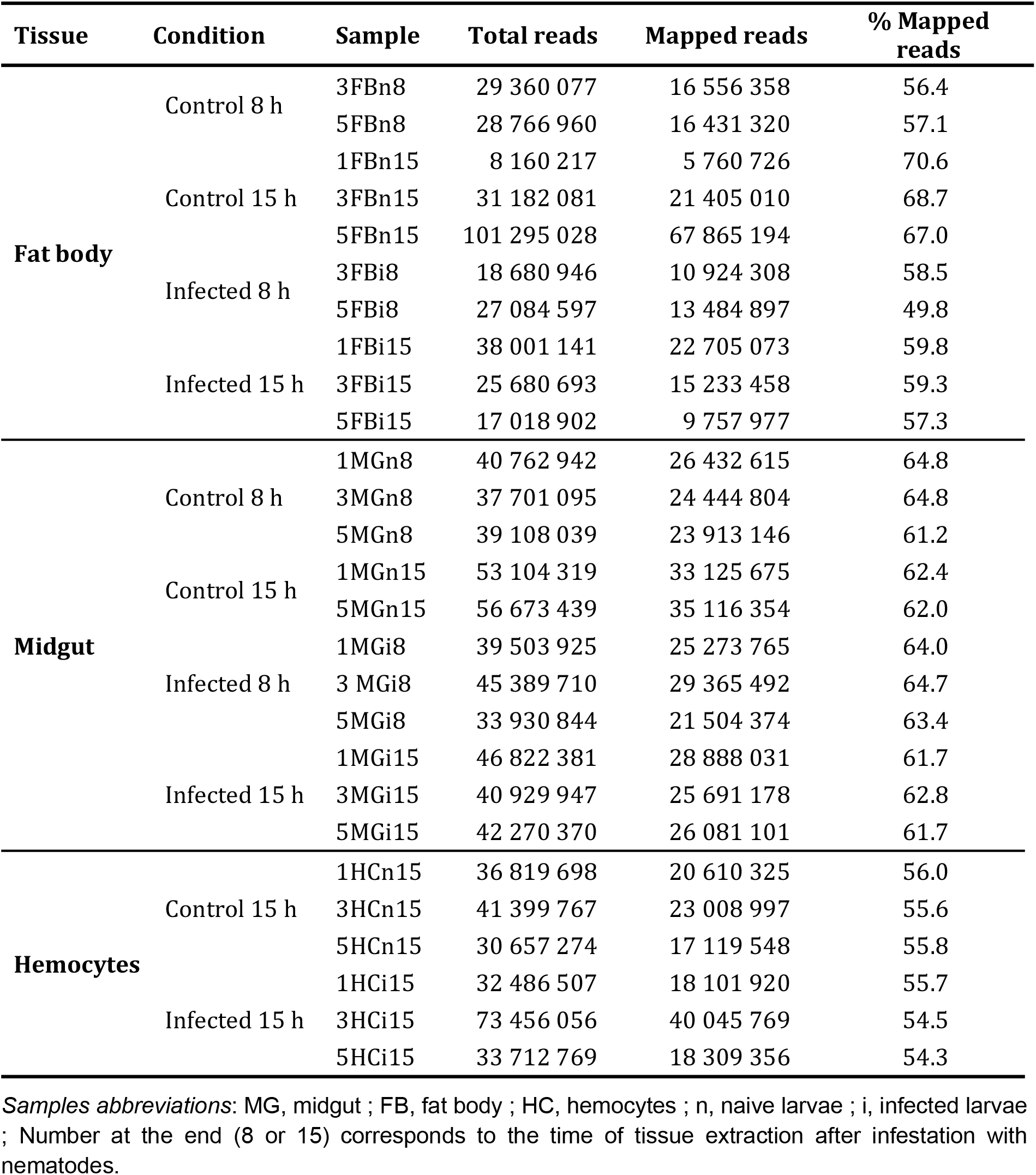
RNAseq statistics

### Overview of transcriptional response

In MG and FB, at 8 hpi, we detected a very small transcriptional response with only 5 statistically significant differentially expressed (DE) genes in FB and none in MG (**S2 Fig**). From these, 4 genes are overexpressed in response to infestation and are also retrieved overexpressed at later time-points in all 3 tissues (**S3 Fig**). They are annotated in the genome as unknown transcripts.

At 15 hpi, there is a more important transcriptional response with thousands of DE genes in all three tissues at padj < 0.01 (**S2 Fig**), with in each case, more genes overexpressed than underexpressed. In order to detect the most significant genes responding to EPN infestation at 15 h, we modeled the EPN effect across all datasets and identified 271 DE genes (Fig 1D) which we intersected with all previous 1 on 1 comparison per conditions (see **Methods**). Of these, we identified a total of 216 DE genes at 15 hpi in the three different tissues (Fig 1E). Most of the response occurred in FB and HC tissues (Fig 1E, Table 2) with, again, a vast majority of overexpressed genes (Fig 1D and 1E, Table 2).

The data obtained by RNAseq were confirmed by quantitative RT-PCR on a selection of DE genes in the three tissues (**S4 Fig**).

**Table 2:** Differentially expressed genes per condition at 15 hpi

| Tissue | Midgut (MG) | Fat body (FB) | Hemocytes (HC) |
| --- | --- | --- | --- |
| **Differentially expressed genes ($\text{padj} < 0.01$)** | 54 | 125 | 189 |
| **Overexpressed genes ($\text{padj} < 0.01$, $\text{Log2FC} > 0$)** | 41 (75.9 %) | 115 (92.0 %) | 182 (96.3 %) |
| **Underexpressed genes ($\text{padj} < 0.01$, $\text{Log2FC} < 0$)** | 13 (24.1 %) | 10 (8.0 %) | 7 (3.7 %) |

### Functional response of the Midgut at 15 hpi

The first line of defense against *S. carpocapsae* EPN is the midgut barrier, which is also the main entry point of *Steinernema* nematodes in *S. frugiperda*. This organ is known for its immune activity through the production of reactive oxygen species and anti-microbial peptides in response to pathogens (27). It is not supposed to be directly confronted to *X. nematophila* and therefore, we hypothesized that the genes, which would be overexpressed specifically in the midgut may identify anti-nematode factors. However, we did not find any DE genes in the midgut at 8 hpi and only 4 genes that were DE specifically in the midgut tissue at 15 hpi, with only 1 being overexpressed and 3 under-expressed (Fig 2A). The overexpressed gene encodes a heat-shock protein of the hsp70 family (**S1 Data**). This superfamily of genes is usually upregulated in response to oxidative stress and in the midgut of *Drosophila*, Hsp68 promotes the proliferation of intestinal stem cells, and thus its regeneration (28).

**Fig 2:**
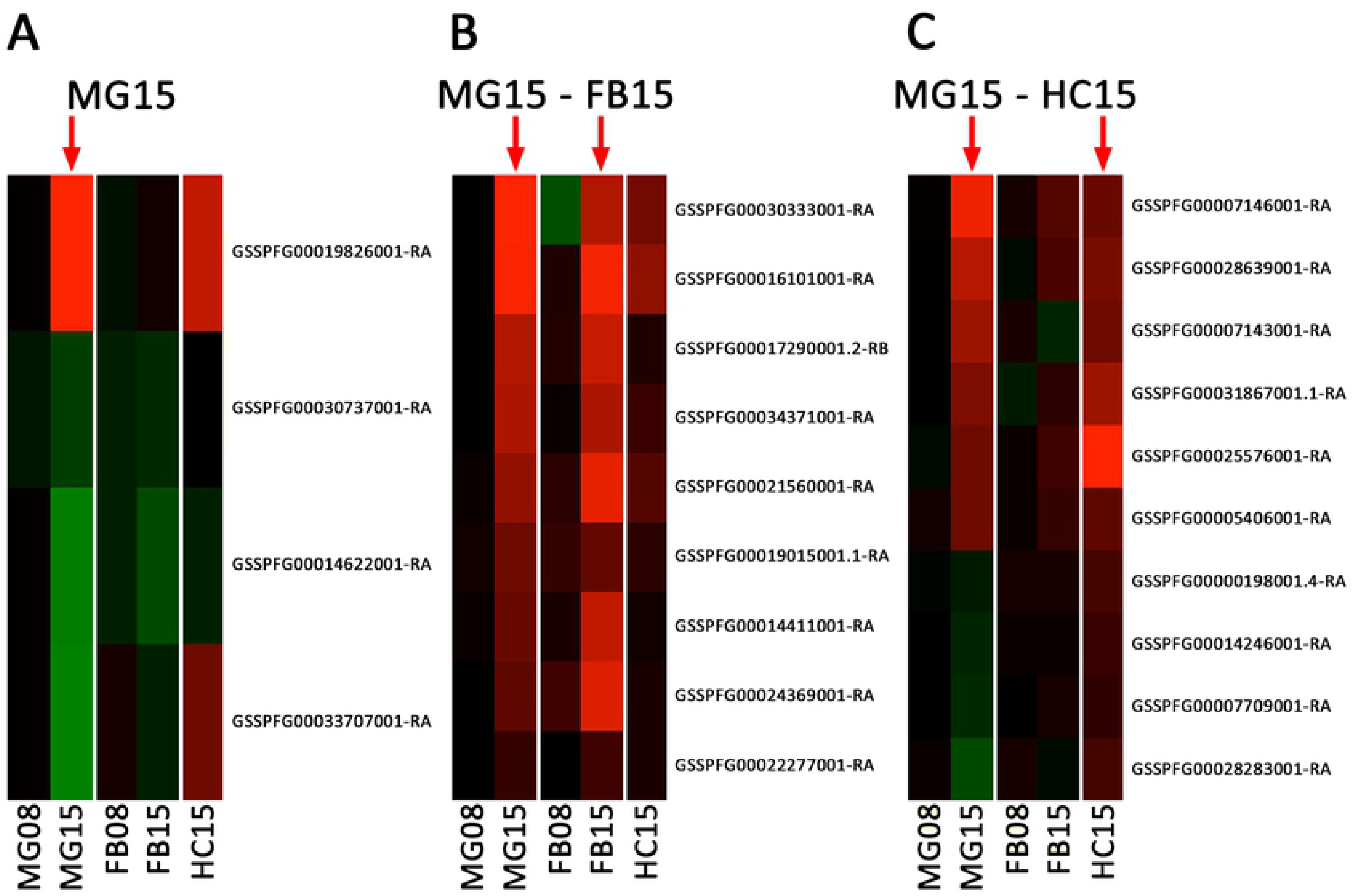
Midgut Associated Response. Heatmaps of differential expression (in log2FoldChange - green, under-expression, red, overexpression according to values in **S1 Data**) across all experimental conditions of genes found significantly differentially expressed **A:** specifically in the MG at 15 hpi (MG15 - 4 genes) **B:** common to MG15 and FB15 (9 genes) and **C:** common to MG15 and HC15 (10 genes).

One of the under-expressed genes is an E3 ubiquitin-protein ligase of the Seven In Absentia family (SIAH). The mammalian homologue Siah1 cooperates with SIP (Siah-interacting protein), the F-box protein Ebi and the adaptor protein Skp1, to target beta-catenin, a multifunctional protein that plays an important role in the transduction of Wnt signals and in the intercellular adhesion by linking the cytoplasmic domain of cadherin, for ubiquitination and degradation via a p53-dependent mechanism. Thus, down-regulation of SIAH might increase levels of ß-catenin, which favors proliferation of intestinal stem cells in *Drosophila* (14).

The two other under-expressed genes are an ABC family transporter and an NT-C2 domain protein, for which no obvious link to infection or to intestinal homeostasis can be established from the literature.

While we found no evidence of a direct response to EPN specifically by the midgut, we investigated whether this tissue may share a common immune response with the fat body or the hemocytes.

We found nine common DE genes between the midgut and the fat body, which are all overexpressed in both tissues (Fig 2B). Of those, three genes have no annotated structure or function. Four other genes with no clear homology have domains that can be associated to regulation (protein-kinase domain, calcium binding domain, amino-acid transporter and MADF domain transcription factor) (**S1 Data**). Interestingly, we found a cytochrome P450 gene encoding the CYP340L16. CYP genes are usually involved in detoxification of foreign chemicals such as plant xenobiotics and pesticides (29). Finally, one trypsin inhibitor-like cysteine rich domain proteinase inhibitor was also overexpressed. There is no enrichment for a specific molecular function or biological process among those nine genes.

Similarly, 10 genes were found significantly differentially expressed in both the midgut and hemocytes, with 6 of them being overexpressed in both tissues and 4 being under-expressed in MG and upregulated in HC (Fig 2C). Of the 6 overexpressed genes, 4 are small solute transporters of the Major Facilitator Superfamily (MFS), 1 is an antennal carboxylesterase and 1 is a proteinase inhibitor (**S1 Data**). No particular function of note has been identified for the 4 genes that were under-expressed in MG but overexpressed in HC, except for Iap2, which is an inhibitor of apoptosis and a member of the IMD pathway (30).

From this comparison, it seems likely that the midgut is not specifically mobilized to defend the *S. frugiperda* larvae against EPN infestation. There are few DE genes in this tissue, whether specific or in common with other tissues, and no specific functional pathway can be clearly identified. Rather, some of the genes identified may be reacting to oxidative stress and homeostasis maintenance of the intestinal epithelium, which might be consequences of the host infestation.

### Specific response of the fat body at 15 hpi

In insects, the adipocytes that compose the fat body are in direct contact with the hemolymph. The physiological function of this organ is to store energetic reserves, in the form of glycogen and lipids, and release them if needed (31). It is also the main tissue involved in systemic immunity, since it produces AMP during immune challenge (8). We found 14 DE genes specifically in the fat body, with 12 overexpressed and 2 under-expressed genes. Among the latest are 1 uncharacterized protein and 1 putative transposable element (TE). Among the 12 fat body-specific up-regulated genes, we found one major actor of immunity, the Toll receptor (32), which recognizes the cleaved circulating cytokine Spätzle to induce the production of anti-microbial peptides (33). It is noteworthy that Toll receptor is expressed in all three tissues, but overexpressed in response to EPN in the fat body only (Fig 3A). Among the 11 other overexpressed genes, we found one potential receptor of the arrestin family, several enzymes (lipase, carboxypeptidase and a GTPase co-factor), again a MADF-domain transcription factor and a dynein (**S1 Data**).

**Fig 3:**
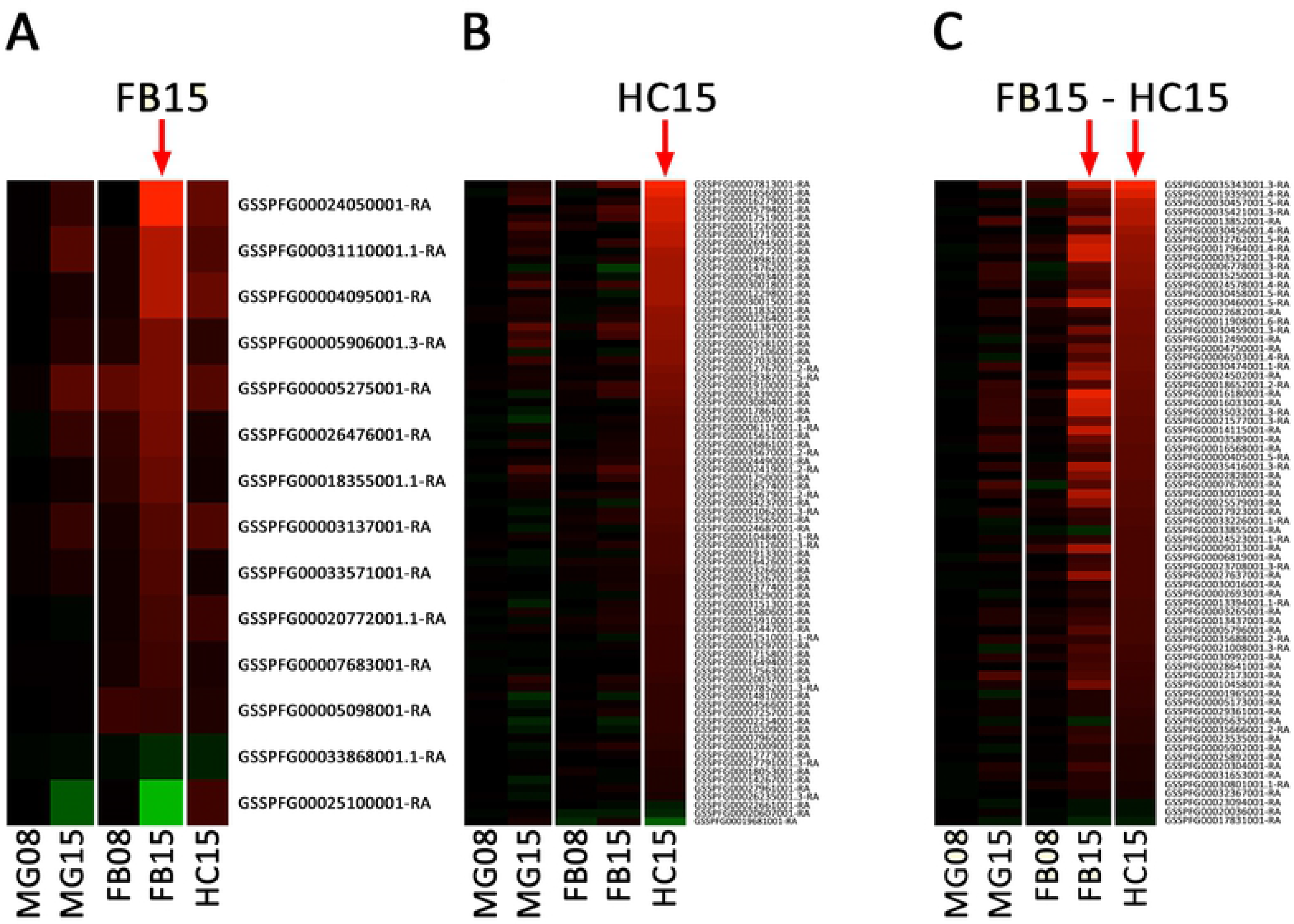
Fat body and hemocytes associated responses. As in Fig 2, heatmaps of differential expression for genes **A:** specific to FB15 (14 genes), **B:** specific to HC15 (77 genes) and C: common to FB15 and HC15 (71 genes).

### Specific response of the hemocytes at 15 hpi

In insects, hemocytes are the main actors of the cellular immune responses like phagocytosis or encapsulation (34). They are also involved in other defense mechanisms such as coagulation (35), and melanization (36, 37). In addition, different reports have shown that hemocytes are, as the fat body, capable of synthesis of AMP (38–40). Therefore, we hypothesized that genes specifically induced in the hemocytes may be involved in coagulation and/or melanization along with cellular immune responses.

The largest number of DE genes is found specifically in the hemocytes with 77 genes (Fig 1E), of which 74 are overexpressed in response to EPN infestation (Fig 3B). No enriched GO categories have been detected in this list. However, we noticed several categories of genes of interest. The most overexpressed gene encodes a serine protease without CLIP-domain (**S1 Data**) that is homologous to hemolymph proteinase 7 (HP7) and 10 (HP10) in *Manduca sexta* (41). In the insect immune system, serine proteases participate in the activation of Toll-dependent response to infection as well as in the PPO-dependent melanization cascade (42). However, many serine-proteases, in particular without CLIP-domains such as HP7 & HP10, still have unknown function. They are regulated by protease inhibitors, a large family of small peptides, one of which is also found highly induced in our list (**S1 Data**) (log2FoldChange = 8.06). This induction suggests a specific role of HP7/HP10 in the hemocytes activation after infestation by EPN.

The molecular functions we encountered in the hemocytes specific gene list include MFS transporters, ubiquitin-conjugating enzymes, sina-like, antennal esterases, heat-shock proteins and several protein kinases. We also noticed several genes that may play a role in vacuolar trafficking and signaling, with several transmembrane domain proteins. The general molecular function of these genes makes it hard to link them to any biological process.

Surprisingly, we noticed that very few genes linked to immunity were present in this list. In particular, we have not found a deregulation of genes linked to the activation of the PPO pathway besides the above-mentioned serine proteases. The only gene that we could relate to melanization is homologous to the L-dopachrome tautomerase Yellow-f2 that is responsible for the conversion of DOPA into dopamine, a precursor of melanin.

Three transcription factors are also found overexpressed in HC, including Vrille. Vrille is known to activate the serine protease Easter that, in turns, cleaves the Spätzle protein that is the ligand of Toll receptor, which we found overexpressed in the fat body (see above). Other genes involved in immunity are Pellino which might be either a negative regulator (43) or an enhancer (44) of the Toll pathway, and IMD which is a member of the IMD pathway.

### Common response of the Fat Body and the Hemocytes at 15 hpi

We identified 71 genes differentially expressed upon EPN infestation in both FB and HC tissues (Fig 1E), 66 of them being overexpressed in both tissues. The “immunity” ontology is the most enriched GO category (**S5 Fig**). The most overexpressed genes correspond to a battery of anti-microbial peptides (attacins, cecropins, defensins, gloverins and moricins) (**S1 Data**).

We identified by homology a repertoire of 40 AMP in the genome of *S. frugiperda*, classified in 7 different families (25). The majority of AMP production is performed by the FB tissue with members of the attacins, cecropins, gloverins and lebocins strongly overexpressed (Fig 3C). These AMP are also significantly overexpressed in the HC but to a lesser extent (Fig. 3C). Defensins such as the gallerimycin (45) and Spod-x-tox (46) are overexpressed in both tissues. Remarkably, of the 10 moricins present in *S. frugiperda* genome, only Moricin 10 is strongly overexpressed in both tissues. The diapausin overexpression is less clear with low levels of expression.

Among the most overexpressed genes in both FB and HC tissues, we also identified several members of the peptidoglycan recognition proteins (PGRP) (**S1 Data**), a family of receptors, which are involved in the recognition of pathogens associated molecular patterns (PAMP) and in the subsequent activation of the Toll, Imd and PPO system pathways (19, 20). In the genome of *S. frugiperda*, we identified 10 PGRP (**S6A Fig**) that were named according to *Bombyx mori* nomenclature (9). Upon EPN infestation, PGRP-S2, -S6 and -L3 are overexpressed in both FB and HC, with PGRP-S2 being the highest overexpressed (**S1 Data**). A phylogenetic analysis (**S6B Fig)** shows that PGRP-S2 is closely related to *Drosophila melanogaster* PGRP-SA, which is involved in the induction of the Toll pathway (47). Recently, it has also been reported that a PGRP-SA homolog may be responsible for the activation of the phenoloxidase system in the Chinese tussar moth *Antheraea pernyi* (48).

This overexpression of AMP and PGRP upon EPN infestation has been similarly observed in transcriptomic studies of *Drosophila melanogaster* larvae infested by *Heterorhabditis* sp. or *Steinernema* sp. nematodes (22–24).

Other immunity genes overexpressed in both tissues are implicated in the Toll pathway. This pathway is activated by PAMP recognition proteins or by proteases from pathogens (21, 49). Signal is transduced by an intracellular complex (MyD88/Tube/Pelle) that binds to the intracytoplasmic domain of Toll and is regulated by Pellino (43, 44). Signal transduction results in the phosphorylation of the ankyrin-repeat containing protein Cactus, which allows its dissociation from the transcription factor Dorsal. This dissociation promotes the translocation of Dorsal to the nucleus where it activates the production of AMP. Several members of this pathway are overexpressed in both FB and HC at 15 hpi, including Pelle, Pellino, and Cactus. Altogether, these results suggest that the Toll pathway is activated in both the fat body and the hemocytes.

We also observed the overexpression of Hdd23 which mediates PPO activation (50) and of 3 putative transcription factors, one of them containing a zinc-finger domain (GATA4-like) known to mediate immune response in *Drosophila* (51).

In addition to immune-related genes, this list contains 7 protease inhibitors. One is a serine protease inhibitor, which may be involved in the regulation of serine proteases cascades, such as the prophenoloxidase activating cascade (42) or the Toll activating cascade (8), while another has homology with a tissue inhibitor of metalloproteases (TIMP). More interestingly, the 5 remaining protease inhibitors belong to the family of inducible metalloprotease inhibitors also called IMPI (52). These 5 IMPI are present as a cluster of genes in the genome of *S. frugiperda* (Fig 4A) and their expression is upregulated especially in the hemocytes (Fig 4B). The first IMPI was purified from the hemolymph of the greater wax moth *Galleria mellonella* (53) and further cloned (54). The expression of IMPI, along with antimicrobial peptides/proteins, is induced by metalloproteases released by damage tissue or metalloproteases from pathogens during the humoral immune response of *G. mellonella* (52). It is well known that entomopathogenic bacteria, such as *X. nematophila* or *Bacillus thuringiensis*, establish their pathogenesis by secreting virulence factors among which metalloproteases (55–59). Therefore, we may hypothesize that *S. frugiperda* induces the expression of IMPI to counteract the metalloproteases produced by *X. nematophila*.

**Fig 4:**
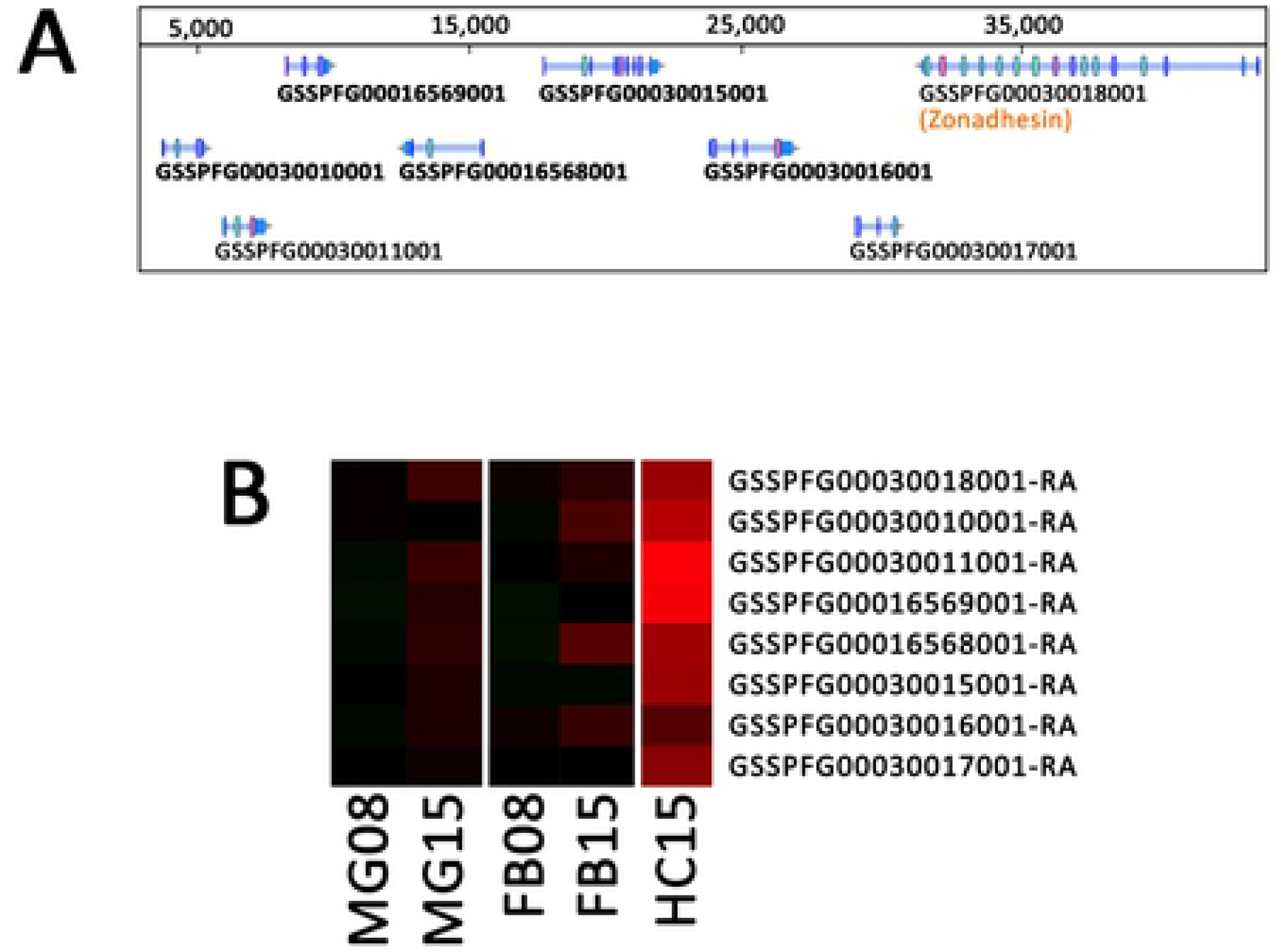
IMPI. **A:** WebApollo viewer showing the annotation of inducible metalloprotease inhibitors genes in cluster on the scaffold_1741 in the genome of *Spodoptera frugiperda*. **B:** As in Fig 2, heatmap of differential expression for the identified IMPI genes.

### Common response of the Midgut, the Fat body and the Hemocytes at 15 hpi

Finally, we identified 31 genes that are significantly differentially expressed in all three tissues (MG, FB and HC), 25 of them being overexpressed in all tissues (Fig 1E**, S1 Data**). There is no enrichment for a specific molecular function or biological process among these 25 genes and only 3 of them could be related to the caterpillar defenses. They encode the previously cited Hdd23 and Cactus, plus Relish, the transcription factor of the Imd pathway, suggesting that this anti-Gram negative bacteria immune pathway (60) could also take part in the previously described humoral responses.

During the annotation of these genes, we noticed that the 4 most differentially expressed genes had no known function. Three of them were also among the few that were overexpressed in the FB at 8 hpi (Fig 5A). We pursued the investigation on the potential origin of these genes, which led us to the identification of 2 previously uncharacterized clusters (Fig 5B **and 5C**).

**Fig 5:**
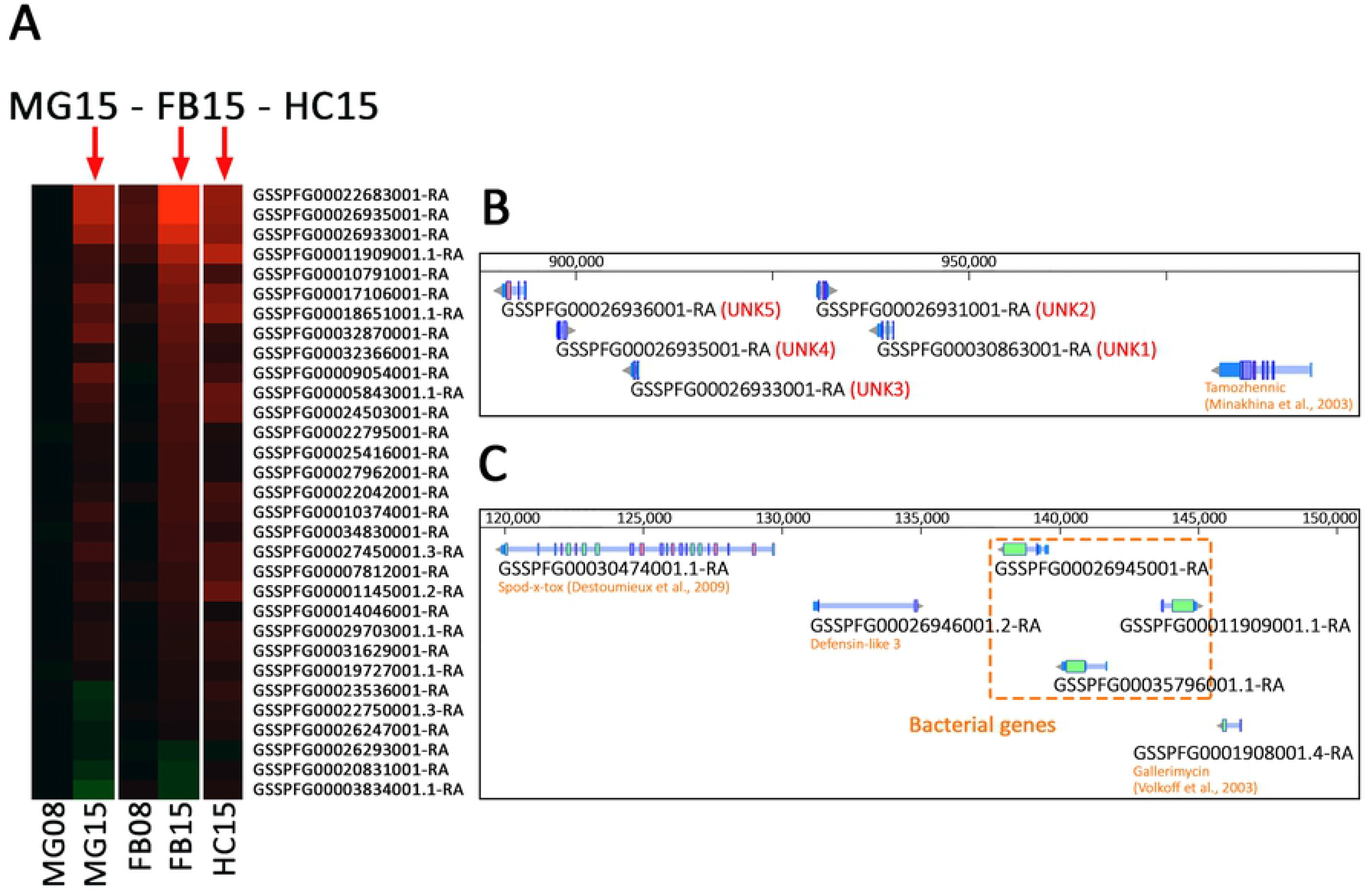
Common tissues. **A:** As in Fig 2 **and** 3, heatmap of differential expression for the 31 genes common to MG15, FB15 and HC15. **B:** WebApollo viewer showing the annotation of unknown genes in cluster on the scaffold_520. **C:** WebApollo viewer showing the annotation of clustered genes of bacterial origin within a defensin cluster.

The first cluster (Fig 5B) is composed of 5 genes for which we could not find any homology in sequence databases at the protein nor at the nucleotide level in any other organism than *Spodoptera frugiperda*. However, we could find the whole cluster in the Sf9 and Sf21 cell lines genomes recently published by other labs (61, 62). In addition, after careful exploration of the syntenic regions, this cluster was also identified in the genomes of two other noctuid species, *Spodoptera litura* and *Helicoverpa armigera* (63, 64). Interestingly, the cluster is located close to a gene homolog to the *D. melanogaster* tamozhennic, which has been reported to be involved in Dorsal nuclear translocation (65). Their genomic localization, their organization in cluster, the presence of eukaryotic signal peptides in their predicted amino acid sequences, the fact that they were not only the most differentially expressed genes upon EPN infestation but also the earliest differentially expressed, led us to the hypothesis that they might encode a new class of immune effectors restricted to some noctuid species.

A second intriguing category of genes in this list has a homology to bacterial proteins of unknown function. They are a set of three genes in cluster (Fig 5C), localized between several defensin encoding genes. They possess a eukaryotic peptide signal and two introns but their main coding sequence is homologous to genes from the *Lactococcus lactis* bacteria (**S7 Fig**) and shares a homology with a cysteine peptidase domain of the papain family. Among insects, these genes are found only in the genomes of other Lepidoptera (**S7 Fig**). Acquisition of antimicrobial activity from bacteria to eukaryotes by horizontal gene transfer (HGT) has been documented before (66), but not in insects. The genes we discovered by transcriptomic here might represent a Lepidoptera specific expansion of immune competence by acquisition of bacterial genes.

## Conclusions

In this work, we have conducted a time-series analysis of tissue-specific transcriptomic response of the Lepidoptera *Spodoptera frugiperda* larvae to the infestation by the EPN complex *Steinernema carpocapsae*/*Xenorhabdus nematophila*. We show that at 8 hours after infestation only a few genes are mobilized in the fat body and none in the midgut, despite the presence and amplification of the EPN symbiont *X. nematophila* in the hemocoel. However, we observed a strong response of the larvae at 15 hours post infestation. This response corresponds to a complementary activation of the immune system by the fat body and the hemocytes, resulting in the production of a large repertoire of humoral effectors and receptors. When we compare our data to RNAseq analyses performed by other laboratories, albeit on different interaction systems, we observe a very similar response. For example, Castillo et al. (2015) report an activation of the immune system of *Drosophila melanogaster* larvae by the EPN complex *Heterorhabditis bacteriophora*/*Photorhabdus luminescens*, represented by several AMPs, several peptidase and protease inhibitors and also Yellow-F. This suggests that all potential hosts possess a conserved ability to fight against EPN complexes.

Despite this powerful response, *S. carpocapsae*/*X. nematophila* complex will be successful regardless of the system they will be confronted with. Anatomies of their host might differ but IJs will find their way inside the midgut and pierce it to enter the hemocoel. There, they will survive the inflammatory response of their host for a sufficient amount of time in order for their released bacterial symbionts to multiply within the insect and kill it (6, 7, 67). It was proposed that the nematodes were able to camouflage themselves from the insect immune system (68). The facts that we found very few genes mobilized at the early time point of 8 hpi and that very few immune-related genes were found mobilized in the midgut at any time point support this idea. However, despite a previous study suggesting that *X. nematophila* could resist to the humoral immune responses by transcriptional down-regulation (69), our study and recent others (24, 70) clearly show that, regardless of the host insect, the immune system is triggered and will react to the EPN infestation.

Nothing in our data suggests a mechanism by which the EPNs bypass the insect defenses at 15 hpi. Indeed, at this time point, all signaling pathways seem activated in both the fat body and the hemocytes and 16 different AMP are produced. Several studies have evidenced a loss of hemolymph AMP and antimicrobial activity during infection by the EPN or by *X. nematophila* (71, 72). A likely hypothesis might be that these AMP are degraded by *X. nematophila* virulence factors, as shown for the *X. nematophila* protease II in *Galleria mellonella* and *Pseudaletia unipuncta* (56).

## Materials and Methods

### Insect rearing

Corn variant *Spodoptera frugiperda* (Lepidoptera: Noctuidae) larvae were reared on a corn-based artificial diet (73). They were maintained at 23°C +/-1°C with a photoperiod of 16 h/8 h (light/dark) and a relative humidity of 40% +/-5%. *Galleria mellonella* (Lepidoptera: pyralidae) larvae were reared on honey and pollen at 28°C in dark.

### Nematode production and storage

Symbiotic *Steinernema carpocapsae* (strain SK27 isolated from Plougastel, France) were renewed on White traps (74) after infestation of the wax moth *Galleria mellonella* last larval stages. They were maintained in aerated Ringer sterile solution with 0.1 % formaldehyde at 8°C for two to four weeks to ensure optimal pathogenicity.

### Infestation

Infestation experiments were processed at 23°C in 12-well culture plates. In each well, one second day sixth instar larva of *S. frugiperda* was placed on a filter paper (Whatman) with artificial corn-based medium. For infested larvae, 150 µL of Ringer sterile solution containing 150 *S. carpocapsae* IJs were introduced in each well. 150 µL of Ringer sterile solution was used for control larvae. To control the nematodes’ efficacy, the survival of 12 larvae in both conditions was monitored for 48 hours in each experiment.

### Xenorhabdus nematophila quantification

The concentration of *X. nematophila* in *S. frugiperda* hemolymph after infestation was estimated by CFU counting on NBTA (nutrient agar supplemented with 25 mg of bromothymol blue per liter and 40 mg of triphenyltetrazolium chloride per liter) with 15 µg/mL of erythromycin. For 3 independent experiments and 3 technical replicates, hemolymph was collected by bleeding of 3 caterpillars in 200 µL PBS buffer supplemented with phenylthiourea. The volumes of hemolymph were then estimated and serial dilutions of the samples were plated. CFU were counted after 48 h incubation at 28°C and CFU numbers were then reported to the estimated hemolymph volumes in order to calculate the bacterial concentrations. The hemolymph of naive caterpillars was also plated to verify the absence of bacterial growth.

### RNA extraction

*Spodoptera frugiperda* larvae were bled and hemolymph was collected in anti-coagulant buffer (75). Hemocytes were recovered by a short centrifugation at 800 g for 1 min at 4°C. The hemocyte pellet was immediately flash-frozen. Then, larvae were dissected and fat bodies and midguts were extracted, rinsed with PBS, flash-frozen with liquid nitrogen in eppendorf tubes and conserved at −80°C until use. After thawing, 1 mL of Trizol (Life technologies) was added and pooled organs were grounded with a TissueLyzer 85210 Rotator (Qiagen) with one stainless steel bead (3 mm diameter) at 30 Hz for 3 min. Grounded tissues were transferred in new eppendorf tubes and left at room temperature for 5 min then 200 µL of chloroform (Interchim) were added. The preparations were homogenized and left at room temperature for 2 min. After a centrifugation at 15,000 g and 4°C for 15 min, the aqueous phase was transferred in new eppendorf tubes. Four hundred µL of 70% ethanol were added and nucleic acid extraction was immediately done with the RNeasy mini kit (Qiagen) according to the manufacturer’s instructions. Contaminating DNA was removed by the use of a Turbo DNA-free^TM^ kit (Life Technologies) according to the manufacturer’s protocol.

RNA yield and preparation purity were analyzed with a Nanodrop 2000 spectrophotometer (Thermo Scientific) by the measure of the ratios A_260_/A_280_ and A_260_/A_230_, respectively. RNA integrity was verified by agarose gel electrophoresis. RNA preparations were then conserved at −80°C.

### Library preparation and Illumina sequencing

Library preparation and RNA sequencing were conducted by MGX GenomiX (IGF, Montpellier, France). Libraries were prepared with the TruSeq Stranded mRNA Sample preparation kit (Illumina). In brief, after a purification step with oligo(dT) magnetic beads, polyadenylated RNAs were chemically fragmented. A first cDNA strand was synthesized with random primers and SuperScript IV Reverse Transcriptase (Life Technologies) and the second strand was then synthesized. After the addition of single adenine nucleotides, indexed adapters were ligated to the cDNA ends. Adapters-ligated cDNAs were then amplified by PCR and libraries were validated on Fragment Analyzer with a Standard Sensitivity NGS kit (Advanced Analytical Technologies, Inc) and quantified by qPCR with a Light Cycler 480 thermal cycler (Roche Molecular diagnostics).

cDNAs were sequenced with the HiSeq 2500 system (Illumina) on 50 base pairs with a single-end protocol. In brief, libraries were equimolary pooled and cDNAs were denatured, diluted to 8 pM and injected in the flow cell. The samples were multiplexed by 6 and a PhiX spike control was used. Clusters were generated with a cluster generation kit (Illumina), cDNAs were sequenced by synthesis. Image analysis and base calling were realized with the HiSeq Control Software (Illumina) and the RTA software (Illumina), respectively. After a demultiplexing step, the sequences quality and the absence of contaminant were verified with the FastQC software and the FastQ Screen software, respectively. Raw data were submitted to a Purity Filter (Illumina) to remove overlapping clusters.

### Alignment and counting

For each sample, the reads were pseudoaligned on the *S. frugiperda* reference transcriptome version OGS2.2 (25) using Bowtie2.2.3 (76). Processing of alignment files (.sam files) into sorted .bam files was performed by samtools view and samtools sort commands (77). Read counts for each gene were obtained using samtools idxstats command.

### Differential expression analysis

Differential expression was analyzed with the R package DESeq2 (26). Treated versus untreated samples of the same tissue + time conditions were analyzed using a classical method. An example is shown in **S2 Data** for the analysis of differential expression in the fat body at 8 hpi identifying 5 DE genes at a p-value adjusted of 0.01, equivalent to 1% false discovery rate (**S2 Fig**).

For the global analysis of the EPN effect, we used the Likelihood Ratio test function of DESeq2 as presented in **S3 Data.** At an adjusted p-value of 0.1, this method identified 271 DE genes associated to EPN treatment. We overlapped this list with the pair-wise comparisons above to define tissue-specific or common responses as shown in Fig 1E. Each sub list of DE genes has been analyzed with Blast2GO Pro software (78) to identify homolog sequences by blastx as well as GO categories. By using the full list of *S. frugiperda* OGS2.2 transcripts as reference (25), the enriched GO terms were identified with a Fisher’s exact test (one-tailed, FDR < 0.05).

Heatmaps were generated using the heatmap.2 function of the gplots R package such as this presented in **S4 Data** that generated Fig 1D.

### qPCR and primers

Differential expression data were verified with control RT-qPCR on selected upregulated and downregulated genes on independently performed EPN infestation experiments. cDNA was synthesized with SuperScript II Reverse Transcriptase (Invitrogen) from 1 µg of RNA sample, according to the manufacturer’s protocol.

The primers (**S1 Table**) were designed with the Primer3Web tool (79). Their efficiency was estimated by using serial dilutions of pooled cDNA samples and their specificity was verified with melting curves analysis. Amplification and melting curves were analyzed with the LightCycler 480 software (Roche Molecular diagnostics) version 1.5.0.

RT-qPCR were carried out in triplicate for each biological sample, with the LightCycler 480 SYBR Green I Master kit (Roche Molecular diagnostics). For each couple of sample and primer, 1.25 µL of sample containing 50 ng/µL of cDNA and 1.75 µL of Master mix containing 0.85 µM of primers were distributed in multiwell plates by the Echo 525 liquid handler (Labcyte). After an enzyme activation step of 95°C for 15 min, the amplification was monitored in the LightCycler 480 (Roche) thermal cycler for 45 cycles of 95°C for 5 s, 60°C for 10 s and 72°C for 15 s.

Crossing points were determined using the Second Derivative Maximum method with the LightCycler 480 software (Roche Molecular diagnostics) version 1.5.0. Relative expression quantifications were then processed with the REST 2009 software (80), using the pairwise fixed randomization test with 2,000 permutations. Targets relative levels were normalized to RpL32 housekeeping gene relative levels and the EF1 gene was used as an internal control.

## Acknowledgments

We thank the quarantine insect platform (PIQ), member of the Vectopole Sud network, for providing the infrastructure needed for pest insect experimentations. We are also grateful to Clotilde Gibard and Gaëtan Clabots for maintaining the insect collections of the DGIMI laboratory in Montpellier.

---

**S1 Table: Primers sequences and genes**

**S1 Data: Differentially Expressed (DE) genes at 15 hpi**

This table presents the genes that are found differentially expressed upon EPN infestation at 15 hpi. The baseMean column represents the DESeq2 normalized mean expression levels of a particular gene across all experiments. The log2FoldChange and padj columns provided are given by the DESeq2 ‘results’ command from pair-wise comparisons between (Midgut: MG, Fat Body: FB, Hemocytes: HC). On the right hand of the table, we present the best blastp homolog of each gene as well as our manual annotation of each gene based on this homology and protein domain analyses or on previous annotation (25)(in bold). Genes have been grouped by tissues where the differential expression false discovery rate (padj) was less than 0.01.

**S2 Data: Script used for the analysis of differential expression in the fat body at 8 hpi.**

**S3 Data: Script for global analysis of the EPN effect.**

**S4 Data: Heatmaps were generated using the heatmap.2 function of the gplots R package such as below that generated** Fig 1D.

**S1 Fig: Dataset quality control**

**A:** Heatmap of pair-wise correlation between all RNAseq samples. Hierarchical clustering of samples shows the grouping of experiments mostly by tissues then by time-point and by condition. **B:** Principal Component analysis of RNAseq samples showing the grouping of experiments mostly by tissue then by time-point and by condition.

**S2 Fig: DESeq2 analysis**

MA plots showing the log2FoldChange in function of mean expression (measured in reads coverage) of total *S. frugiperda* transcripts in a pairwise DESeq2 analysis of EPN vs PBS in every Tissue*Time-point conditions. Indicated on each plot is the number of genes significantly overexpressed (red box) or under-expressed (blue box) upon EPN infestation. For experiments at 15 hpi, the total number of DE genes is also indicated on the right side of the plot.

**S3 Fig: DE genes in fat body at 8 hpi**

At 8 hpi, 5 genes are significantly differentially expressed in the fat body (FB08). This heatmap of log2FoldChange shows that the 4 unknown genes significantly overexpressed in the FB08 condition are also overexpressed at 15 hpi in all 3 tissues. On the right hand side, this table indicates the DESeq2 results for these genes, showing that the 4 overexpressed genes are of unknown function and correspond to the gene cluster presented in Fig 5B.

**S4 Fig: Validation of RNAseq data using quantitative RT-PCR**

Selected genes were analyzed by quantitative PCR using RNA samples from tissues of naïve or infected larvae. The relative expression level (ratio infected/naïve larvae) is shown as log2FoldChange mean from 3 independent experiments.

**S5 Fig: Fat Body and Hemocytes Gene Ontology enrichment**

The DE genes common to FB and HC at 15 hpi (Fig 3C) show an enrichment of Gene Ontology terms associated to immunity. The same analysis conducted on the other gene lists, in particular the 77 genes associated to HC15, did not produce significant GO term enrichments.

**S6 Fig: Structures and phylogeny of *Spodoptera frugiperda* peptidoglycan recognition proteins**

**A:** Structures of *Sf*PGRP. In red, signal peptide and in blue, transmembrane domain. **B:** The phylogeny of 95 PGRP domain amino acid sequences was determined by using the Maximum Likelihood method (81) in MEGA7 (82). The bootstrap consensus tree was built from 100 replicates and branches corresponding to partitions reproduced in less than 50% bootstrap replicates were collapsed. Initial tree(s) for the heuristic search were obtained automatically by applying Neighbor-Join and BioNJ algorithms to a matrix of pairwise distances estimated using a JTT model, and then selecting the topology with superior log likelihood value. Branches colors: green, Lepidopteran (Bm: *Bombyx mori*, Dp: *Danaus plexippus*, Hm: *Heliconius melpomene*, Px: *Papillio xuthus*, Sf: *Spodoptera frugiperda*), bleu, Dipteran (Aa: *Aedes aegypti*, Ag: *Anopheles gambiae*, Dm: *Drosophila melanogaster*) and orange, Hymenopteran (Am: *Apis mellifera*, Nv: *Nasonia vitripennis*).

**S7 Fig: Phylogenetic analysis of genes from bacterial origin**

The 100 first best hits after blastp on nr NCBI were retrieved and the phylogenetic tree was constructed using the method described as in **S6 Fig**. Sequences are grouped in two main clades, one with bacteria only and the second with Lepidoptera only.

